# A Non-Intrusive Computer Vision Framework for Real-Time Vital Sign Digitization and Adaptive Drug Infusion in Critical Care Environments

**DOI:** 10.1101/2025.10.17.683010

**Authors:** Anshuli Chikhale, Ninad Mehendale

**Affiliations:** K. J. Somaiya School of Engineering, Somaiya Vidyavihar University, Vidyavihar, Mumbai, 400077, Maharashtra, India

**Keywords:** Computer vision, Vital sign digitization, Adaptive drug infusion, Optical character recognition, Fuzzy logic control, Intensive care automation

## Abstract

**Purpose:** This work proposes a computer vision framework to automate the extraction of vital signs from bedside monitor systems and facilitate adaptive drug infusion in intensive care units (ICUs). This approach is intended to meet the requirement for less manual intervention and improved accuracy in critical care settings.

**Methods:** An 8-megapixel camera captures time-lapse images of a simulated monitor display at a rate of 3 frames per second. Images are preprocessed (grayscale conversion, histogram stretching, edge detection, color filtering) before Tesseract OCR reads out vital signs (heart rate, blood pressure). On a synthetic dataset of 3,000 vital sign images, the system achieved 99.87% accuracy. Retrieved information inputs a pre-programmed fuzzy logic controller to adjust syringe pump infusion rates of medications like norepinephrine. The system is designed for non-invasive integration with existing monitors.

**Results:** OCR accuracy improved from 95.43% (without preprocessing) to 99.87% on the test dataset. An optimal focal distance of 11.9 cm was determined, balancing accuracy and compactness. The fuzzy logic controller provided stepwise adjustment of flow rate based on blood pressure thresholds, with simulations using non-biological substitutes demonstrating predictable control around a 3 ml/hour baseline.

**Conclusion:** This framework offers a promising proof-of-concept for ICU automation, with potential extensions to other monitoring areas. The primary limitation is the lack of human trials and validation in a clinical environment, implying the necessity for future clinical validation before any real-world application can be considered.

## 1 Introduction

Continual monitoring of vital signs and accurate administration of medications are essential in intensive care units (ICUs) to ensure patient outcomes. Conventional manual methods are prone to errors, delays, and increased workload for caregivers. Automation with the help of advanced technology can reduce these drawbacks, improving efficiency and safety.

Computer vision has been a success in medicine for image analysis and patient monitoring. However, its extension to the real-time extraction of vital signs from legacy bedside monitors for direct integration with automated drug delivery systems remains an area with potential for further development. In this paper, we introduce a non-invasive computer vision framework that captures screen-projected vital signs and adjusts drug infusion rates accordingly in a simulated environment.

The system employs a high-resolution camera to acquire images of monitor displays, processes them through optical character recognition (OCR), and uses fuzzy logic to control a syringe pump. This retrofittable approach avoids hardware modifications to the existing medical devices, potentially reducing costs and compatibility issues. Beyond ICUs, the flexibility of the framework could be applicable to other domains such as environmental monitoring or public information systems.

By facilitating real-time processing of data and reactive treatment, this proof-of-concept system aims to demonstrate a pathway to transform critical care and allow clinicians to focus on more intricate tasks with fewer mistakes.

Figure 1 shows the assembly of the system on a vital sign monitor, with a Raspberry Pi managing the camera input and stepper motor-driven syringe output.

**Fig. 1.**
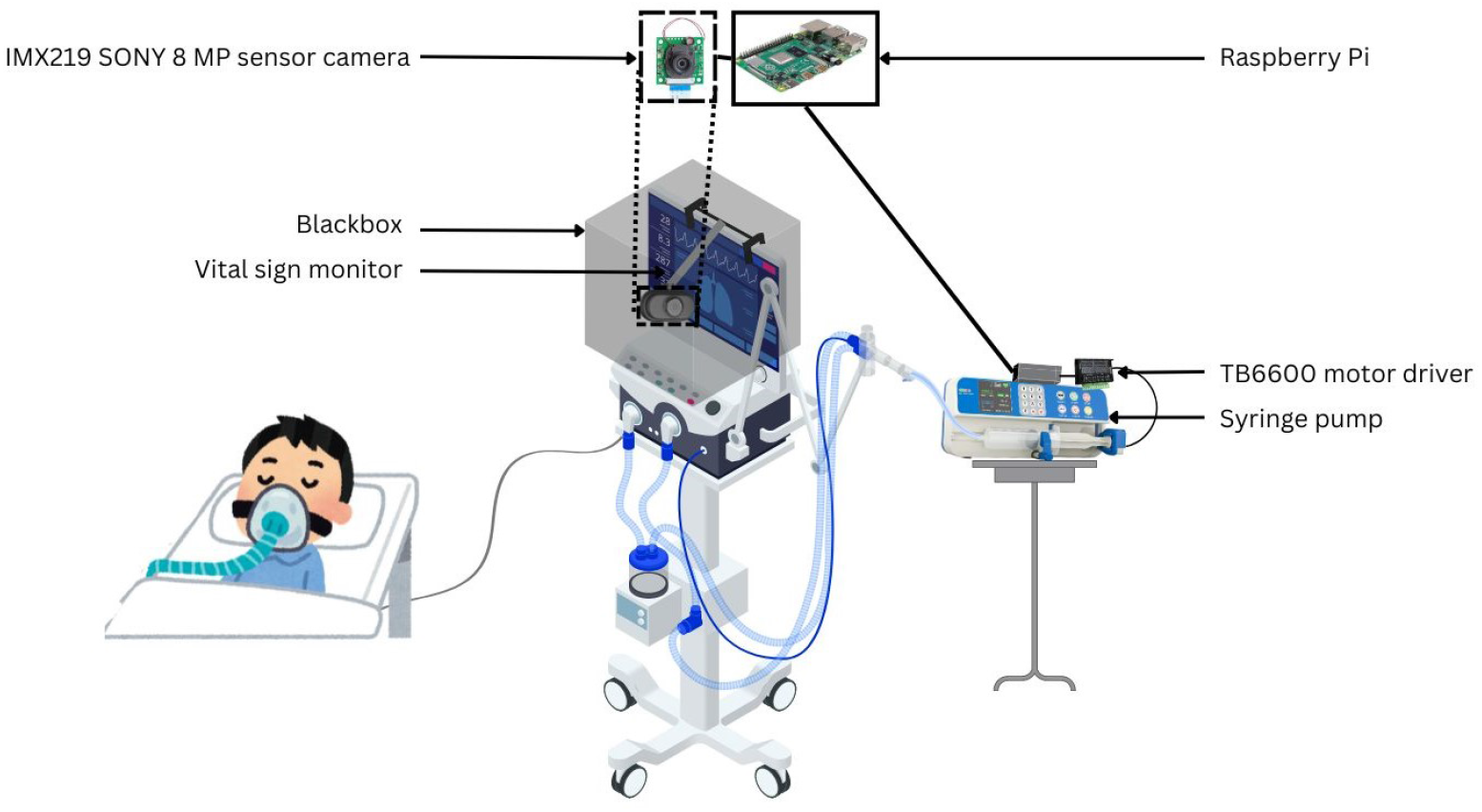
Schematic of a computer vision based system assembly on a vital sign monitor

## 2 Literature Review

### 2.1 Nursing Documentation and Health Information Technology

Computerized healthcare systems try to decrease workload and increase outcome. De Groot et al. (2022) demonstrate in a mixed-methods study that high-quality nursing documentation by community nurses reduces perceived workload, and there are implications for the ICU environment [1]. Shekelle et al. (2006) present an evidence report that health information technology is cost-effective in the long run, and it guarantees economic sustainability of such systems in critical care [2].

### 2.2 Computer Vision and Optical Character Recognition in ICU Monitoring

Vision-based technologies are revolutionizing ICU monitoring. ROMI, a real-time optical digit recognition system utilizing camera-equipped robots to detect ICU equipment, was created by Jeon et al. (2023) to counter the shortcomings of human observations [3]. Froese et al. (2021) suggest an AI-based strategy of continuous bedside data extraction to promote greater clinical data documentation [4]. Boyko et al. (2000) refer to the U.S. Department of Veterans Affairs system as a model for an epidemiological research resource, which can assist vision-based tools [5].

### 2.3 Automated Drug Delivery Systems

Accuracy in drug delivery is most important. Zhang et al. (2022) introduce a microcomputer-controlled drug delivery vehicle in one chip [6], and Moberg (2022) creates a temperature control device for precise administration [7]. Novak Jr (2018) and O’Connor et al. (2021) introduce autonomous and ventilator-based drug delivery systems, respectively, for increased flexibility in critical care [8, 9].

### 2.4 Real-Time and Distant Monitoring of Vital Signs

Non-contact monitoring is essential to ICU safety. Behar et al. (2020) describe non-contact vital sign monitoring during COVID-19 with reduced contact risk [10]. Becking-Verhaar et al. (2023) report high nurse satisfaction with wireless monitoring on general wards with potential scalability [11]. Jorge et al. (2022) demonstrate non-contact monitoring in post-operative ICU patients for the detection of early complications [12].

### 2.5 Regulatory Frameworks and Artificial Intelligence

AI facilitates healthcare automation. Hashimoto et al. (2020) cite AI usage in anesthesiology, for instance, computerized IV anesthesia [13]. The U.S. Food and Drug Administration (2022) gives guidelines for AI/ML-based medical devices [14]. Shaik et al. (2023) describe remote monitoring through AI, citing issues such as data privacy [15]. The conclusions of these studies confirm the model proposed, marking the shift towards automation and clinical validation as central.

## 3 Methodology

The proposed framework employs computer vision techniques and an intelligent control system to enable the automation of critical care environments. The methodological framework, which consists of image processing algorithms, optical character recognition, and a fuzzy inference system to enable extraction of critical signs and adaptive medication dispensing, is described in this section.

### 3.1 Image Acquisition

High-fidelity image capture is the system’s core, employing an IMX219 Sony 8megapixel sensor module with 3280×2464 pixel resolution and a fixed M12 lens mount. The high resolution was chosen to ensure legibility of various font sizes across different monitor models. The camera is deliberately placed at an optimal focus distance of 11.9 cm from the bedside monitor screen, optimized by experimental adjustment for best sharpness (as analyzed in Figure 4). In order to reduce external lighting artifacts, the acquisition system is placed in a light-shielding enclosure. Images are shot in time-lapse with a 3 frames per second (fps) frame rate, synchronized with the OCR processing time. This frame rate was chosen to balance computational overhead with real-time responsiveness, as reflected in the analysis in Figure 5. The acquisition process can be formalized as:

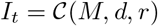

where *I*_*t*_ denotes the image at timestamp *t, C* represents the camera capture function, *M* is the monitor display, *d* = 11.9 cm is the focal distance, and *r* = 3 fps is the capture rate.

### 3.2 Image Preprocessing

Before recognition, images are processed through a multi-stage preprocessing pipeline. The pipeline starts with RGB to grayscale conversion:

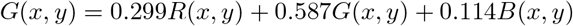

where *G*(*x, y*) is the grayscale intensity at pixel (*x, y*), and *R, G, B* are the color channels. Subsequently, histogram stretching normalizes contrast:

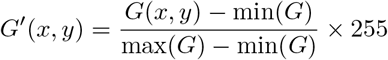

Edge detection employs the Canny algorithm:

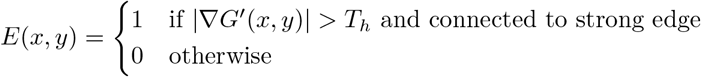

with empirically determined dual thresholds (*T*_*l*_ = 50, *T*_*h*_ = 150) for hysteresis. Finally, color filtering overlays detected edges to accentuate vital sign digits. The complete preprocessing algorithm is outlined in Algorithm 1. Figure 2 illustrates the system workflow, where the camera and Raspberry Pi acquire monitor images, which are preprocessed and passed through Tesseract OCR for vital sign extraction. The recognized values undergo fuzzy logic–based comparative analysis to regulate the syringe pump infusion rate.

#### Algorithm 1

Image Preprocessing Pipeline

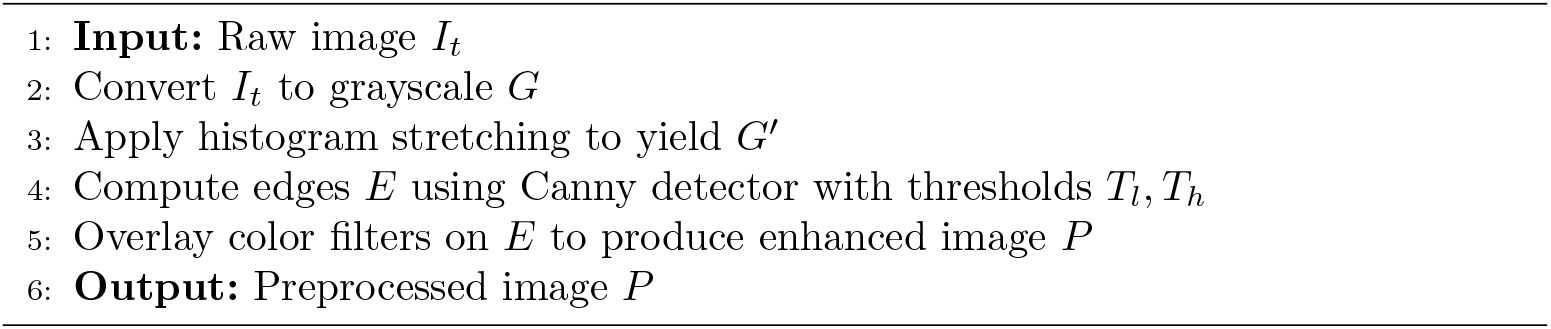

**Fig. 2.**
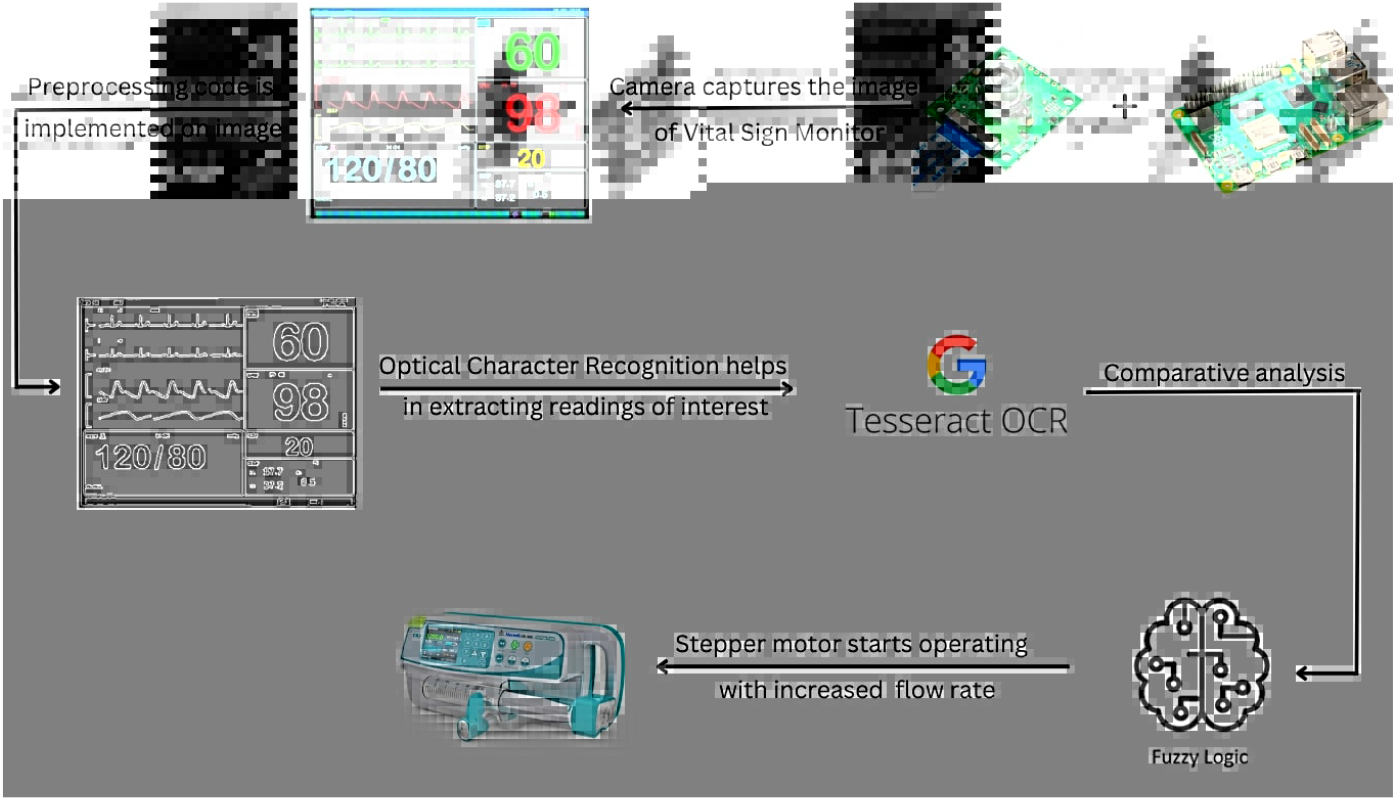
Schematic of the computer vision-based system workflow, highlighting the Raspberry Pi integration with camera and syringe pump.

### 3.3 Optical Character Recognition and Performance Evaluation

The preprocessed images are subjected to Tesseract OCR for digitizing vital signs. To optimize performance, the system crops specific regions of interest (ROIs) where vital parameters are displayed:

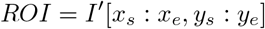

where *I*^*′*^ is the preprocessed image, and (*x*_*s*_, *y*_*s*_, *x*_*e*_, *y*_*e*_) define the bounding box. Tesseract employs LSTM-based recognition. To validate its performance, a synthetic dataset of 3,000 images was generated. These images mimicked the font and color of a common ICU monitor display under varying simulated lighting conditions. Each image contained three numerical vital signs (HR, SBP, DBP). Accuracy was calculated as the percentage of individual numerical values correctly recognized across the entire dataset. The system yielded an accuracy of 99.87% post-preprocessing on this dataset.

### 3.4 Fuzzy Logic Controller

The extracted vital signs serve as inputs to a fuzzy inference system that calculates the appropriate drug infusion rate adjustment. The inputs are systolic blood pressure (SBP), diastolic blood pressure (DBP), and heart rate (HR). These crisp inputs are converted into linguistic categories using membership functions, as defined in Table 1.

**Table 1.** Illustrative Fuzzy Sets for Blood Pressure Inputs.

| Linguistic Variable | Fuzzy Sets (Example Range, mmHg) |
| --- | --- |
| SBP | Low (< 100) |
|  | Normal (100 - 140) |
|  | High (> 140) |
| DBP | Low (< 60) |
|  | Normal (60 - 90) |
|  | High (> 90) |

A rule base, shown in Table 2, combines these linguistic variables to determine the required change in the infusion rate.

**Table 2.** Illustrative Fuzzy Rule Base.

| IF SBP is... | AND DBP is... | THEN Infusion Change is... |
| --- | --- | --- |
| High | High | Strong Increase |
| High | Normal | Moderate Increase |
| Normal | Normal | No Change |
| Normal | Low | Moderate Decrease |
| Low | Low | Strong Decrease |
| ... | ... | ... |

The resulting final drug flow rate is derived through defuzzification using the centroid method:

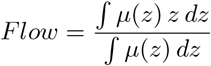

where *μ*(*z*) represents the aggregated output membership. The decision-making workflow is summarized in the flowchart shown in Figure 3.

**Fig. 3.**
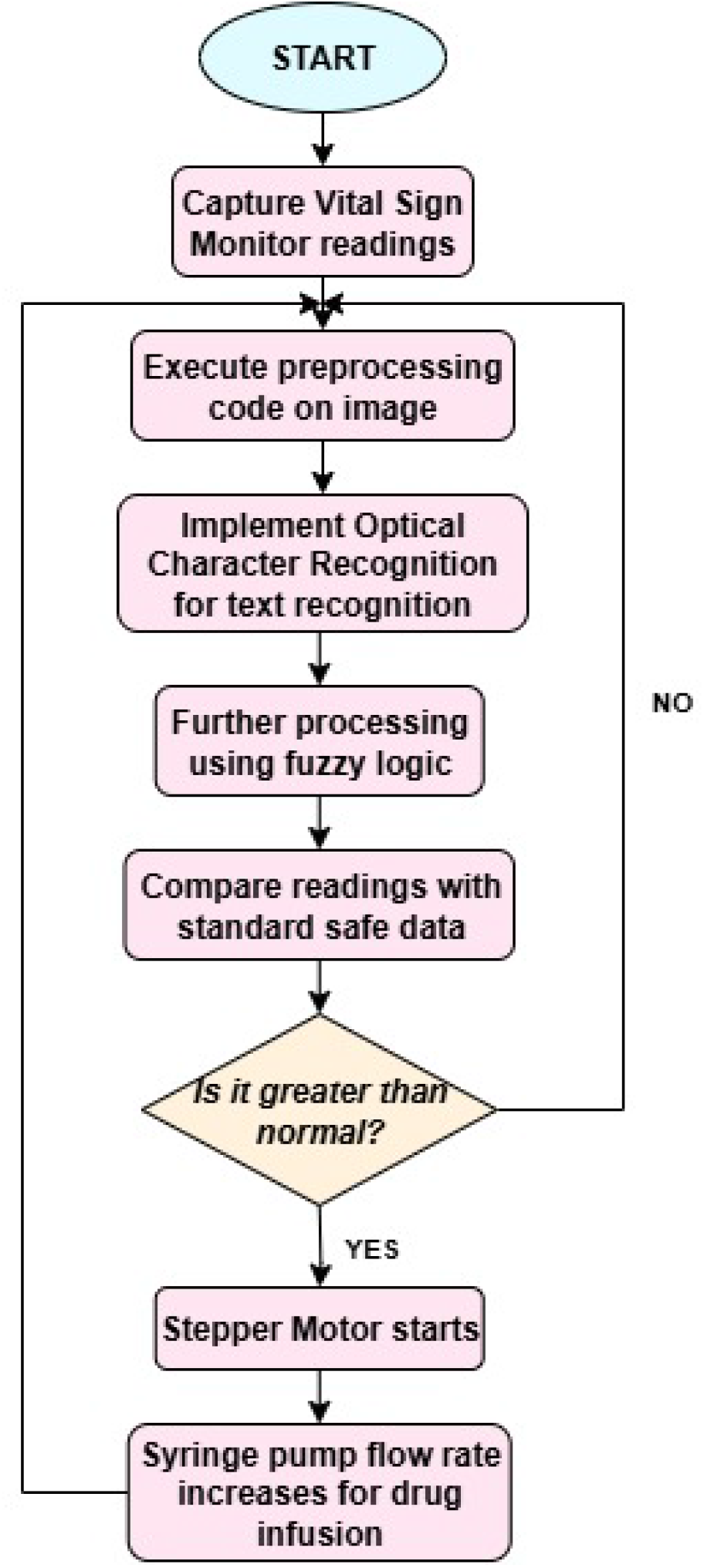
Flowchart of the computer vision-based system for automated vital sign monitoring and drug delivery in intensive care units.

### 3.5 Syringe Pump Control

The computed flow rate modulates a NEMA17 stepper motor via a TB6600 driver. Baseline infusion is set at 3 ml/hour for the simulation, adjusted dynamically for medications like norepinephrine. The control signal is generated as:

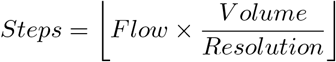

where *Volume* is the syringe capacity and *Resolution* is steps per ml. The system ensures precise delivery by synchronizing motor pulses with the calculated infusion profile. It is important to note this is an open-loop control system; future work would require integrating feedback sensors to correct for mechanical tolerances.

## 4 Results

This section presents experimental outcomes from simulations based on the established methodologies. These results highlight the system’s precision, efficiency, and control capabilities, providing a technical basis for its potential deployment in health monitoring systems.

### 4.1 Focal Distance Optimization and Accuracy Stability

The OCR accuracy of the system demonstrated a clear dependence on the focal distance, as illustrated in Figure 4. The experimental results on our synthetic dataset registered a steep increase from 85.63% at 6 cm to 99.87% at 11.9 cm, leveling off to near 100% beyond 12 cm. The distance of 11.9 cm was selected due to its high accuracy and device compactness (¡ 15 cm depth). This effectively mitigates glare and distortion observed at closer ranges.

**Fig. 4.**
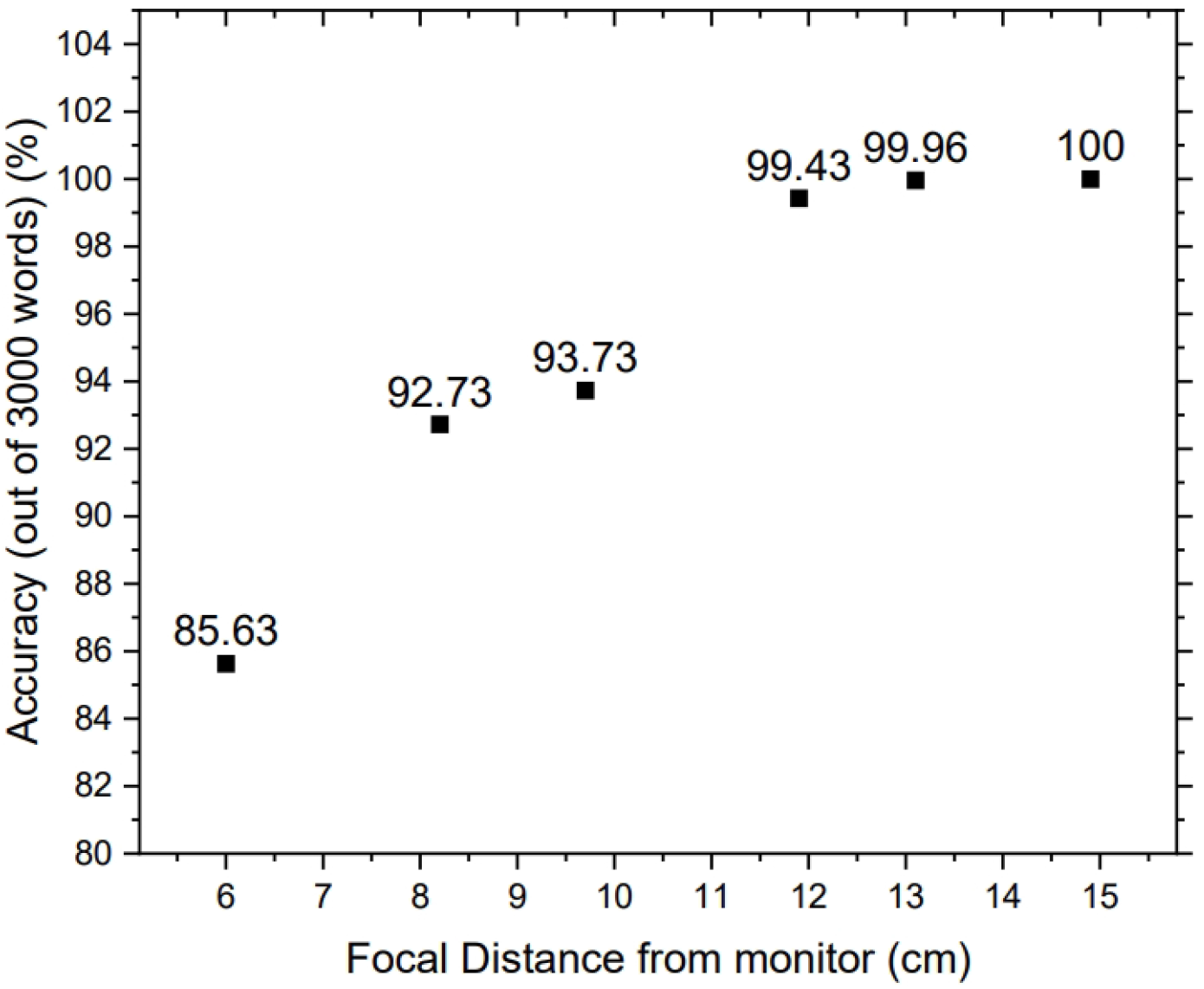
Study of focal distance with respect to OCR accuracy on the synthetic test dataset.

### 4.2 Frame Processing Efficiency and Data Throughput

System throughput was contrasted against different frame rates, as shown in Figure 5, by measuring the number of unprocessed frames in the system’s buffer (SBUF). While operating at 3 FPS, there were zero unprocessed frames, matching the monitor’s 3.85 FPS refresh cycle and providing real-time data availability. At rates greater than 3 FPS, unprocessed frames grew quadratically to 27 at 6 FPS, indicating a processing bottleneck. A regression model (*R*^2^ = 0.98) confirmed an upper limit of sustainable throughput of approximately 3.2 FPS.

**Fig. 5.**
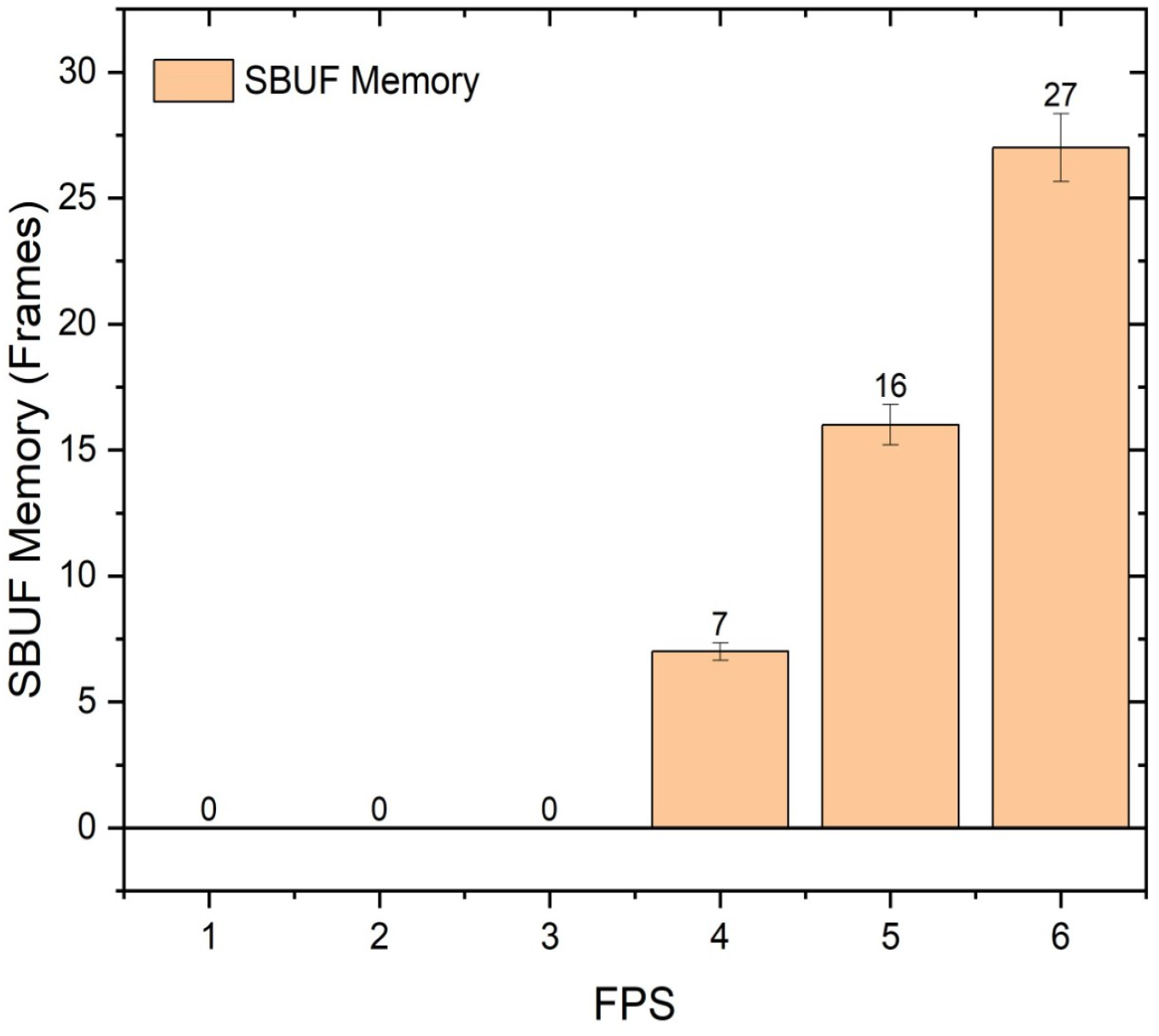
Depiction of unprocessed frames in the system buffer (SBUF memory) versus input camera FPS.

### 4.3 Systolic Blood Pressure-Driven Infusion Modulation

Adaptive infusion control based on systolic blood pressure (SBP) was optimized using the fuzzy logic controller. For the purpose of this simulation, a control logic was designed based on general principles of vasopressor administration. **These thresholds, shown in Figure 6, are illustrative for the model and would require calibration and validation based on specific clinical protocols and patient conditions.** The normalized flow rates rose stepwise: for *SBP <* 120 mmHg, 3%; for SBP 120-129 mmHg, 8%; for SBP 130-139 mmHg, 29%; for SBP 140-160 mmHg, 56%; for SBP 160-180 mmHg, 70%; and for *SBP >* 180 mmHg, 92%. Simulations with non-biological substitutes demonstrated stable adjustments across these ranges.

**Fig. 6.**
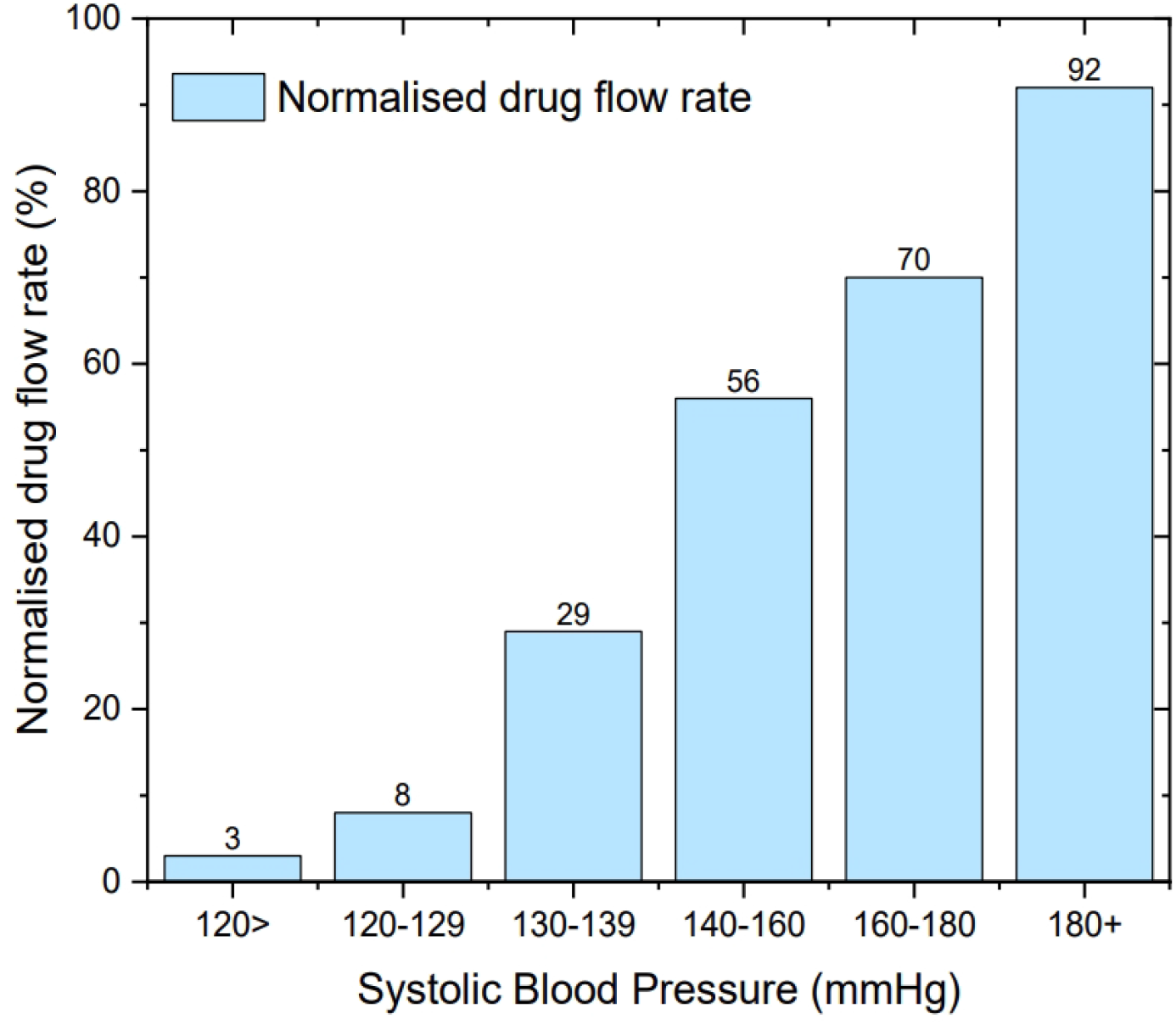
Evaluation of normalized drug flow rate according to systolic blood pressure ranges in the simulation.

### 4.4 Diastolic Blood Pressure-Regulated Safety Thresholds

Diastolic Blood Pressure (DBP) based infusion adjustments yielded an inverse control, as shown in Figure 7. Flow rates reduced from 100% at *DBP >* 180 mmHg to 0% for *<* 60 mmHg to simulate counteracting hypotension risks. Simulations confirmed consistent adherence to these thresholds, highlighting the system’s adaptability to dynamic conditions within the simulated environment.

**Fig. 7.**
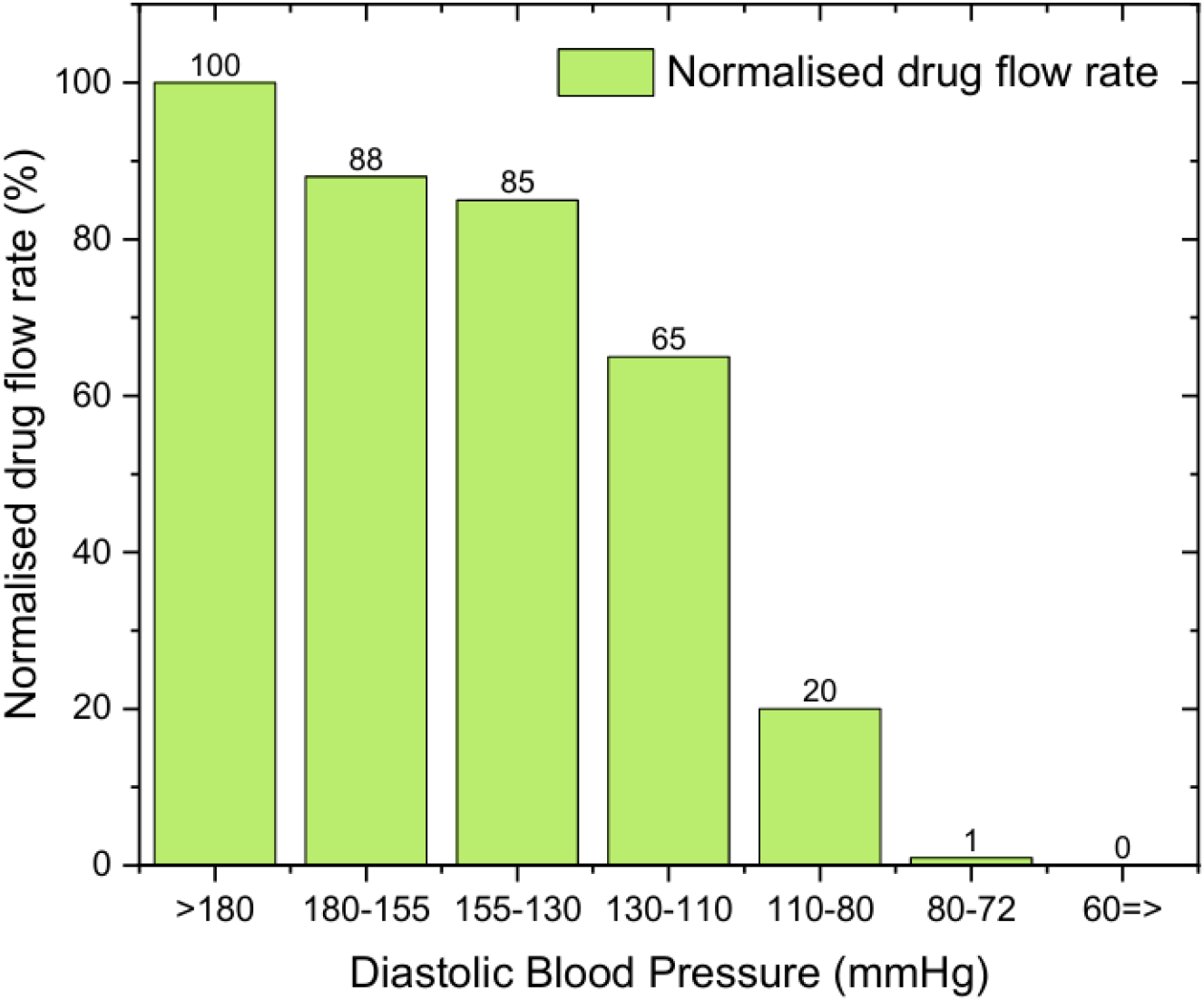
Evaluation of normalized drug flow rate according to diastolic blood pressure ranges in the simulation.

## 5 Discussion

This work introduces a non-intrusive computer vision system that demonstrates the feasibility of improving ICU vital sign monitoring and adaptive drug infusion. The achieved OCR accuracy of 99.87% on our synthetic dataset is promising. The 11.9 cm optimized focal length minimizes trade-offs between image quality and portability. This adds to existing work by Jeon et al. (2023) and Froese et al. (2021), who prioritized real-time data retrieval. A direct comparison of OCR accuracy is challenging as their studies do not report performance on a comparable, standardized benchmark.

The system’s steady-state processing rate of 3 FPS is well-aligned with clinical monitor refresh cycles. The throughput ceiling near 3.2 FPS is a function of computational limits mainly caused by image preprocessing. Breaking such a bottleneck through GPU acceleration or algorithmic improvement offers a path to greater scalability.

The fuzzy logic controller demonstrates the capability to modulate drug infusion rates. The use of linguistic rules to convert blood pressure intervals to dosing changes is in line with recent advances in soft computing for critical care automation.

### 5.1 Limitations and Future Work

While these technological advances are promising, there remain significant limitations.

1. **Simulation-Only Study:** The primary limitation is that this study is based entirely on simulation and surrogate equipment. It has not been tested on real patients or in a clinical environment.
2. **Hardware Practicality:** The use of a light-shielding enclosure may be impractical in a dynamic ICU setting.
3. **Lack of Robustness Testing:** The system has not been tested against a wide variety of monitor brands, fonts, colors, or under adverse conditions like screen glare, partial occlusion, or display of error messages.
4. **Safety:** The current system is open-loop and lacks essential safety features, such as alarms, fail-safes, or the ability to detect sensor disconnection.
5. **Clinical Integration:** Integration with hospital digital environments (e.g., using HL7/FHIR standards) has not been assessed.

Future research must prioritize clinical validation through IRB-approved patient trials to verify system stability and therapeutic effectiveness. Further work should focus on developing robust error correction, fusing data with other physiological sensors, and implementing critical safety interlocks before the system can be considered for any clinical application.

## 6 Conclusion

This work introduces a proof-of-concept for a computerized vital sign digitization and adaptive drug infusion system for ICUs. On a synthetic dataset, it attains a high OCR accuracy of 99.87% with real-time processing at 3 FPS. The combination of an optimized imaging pipeline and a fuzzy logic controller demonstrates a feasible method for infusion control responsive to dynamic blood pressure changes in a simulated setting.

While computational demand currently limits scaling past 3.2 FPS, viable optimizations exist. The non-invasive, retrofit-compatible design of the framework allows for broad potential clinical usefulness without hardware modification. However, subject to rigorous clinical verification and integration into hospital information systems, this method only represents an early step. Much further research is required to explore its potential to enhance patient safety and reduce caregiver burden through precise and smart drug delivery.

## Declarations

### Ethics approval and consent to participate

Not applicable. This study was conducted entirely using computer simulations and laboratory equipment. No human or animal subjects were involved.

### Consent for publication

Not applicable.

### Data availability

The synthetic dataset generated for this study can be made available upon reasonable request to the corresponding author.

### Materials availability

All materials belong to the authors and cannot be sold to anyone.

### Code availability

Code will be made available upon reasonable request to the corresponding author.

### Author contribution

Anshuli Chikhale (AC) and Ninad Mehendale (NM) conceptualized the work. AC conducted literature review, collected data, and performed formal analysis. AC and NM were responsible for hardware building and program ming. AC drafted the manuscript. NM reviewed and edited the manuscript. AC and NMcontributed to graphic designs and figures.

